# Silencing of E-cadherin in induced human pluripotent stem cells promotes extraembryonic fates accompanying multilineage differentiation

**DOI:** 10.1101/2020.11.01.363713

**Authors:** Ashley RG Libby, Ivana Vasic, David A Joy, Martina Z Krakora, Fredrico N Mendoza-Camacho, Bruce R Conklin, Todd C McDevitt

## Abstract

In embryonic development, symmetry breaking events and the mechanical milieus in which they occur coordinate the specification of separate cell lineages. Here, we use 3D aggregates of human pluripotent stem cells (hPSCs) encapsulated in alginate microbeads to model the early blastocyst prior to zona pellucida hatching. We demonstrate that 3D confinement combined with modulation of cell-cell adhesions is sufficient to drive differentiation and collective migration reminiscent of the pre-implantation embryo. Knockdown of the cell adhesion protein CDH1 in encapsulated hPSC aggregates resulted in protrusion morphologies and emergence of extra-embryonic lineages, whereas unencapsulated CDH1(-) aggregates displayed organized radial delamination and mesendoderm specification bias. Transcriptomic similarities between single-cell RNA-sequencing data of early human embryos and encapsulated CDH1(-) aggregates establishes this *in vitro* system as a competent surrogate for studying early embryonic fate decisions and highlights the relationship between cell-cell adhesions and the mechanical microenvironment in directing cell fate and behavior.

**Highlights:**

- Generation of embryonic scale 3D morphogenesis using hydrogel encapsulation
- Manipulating adhesion triggers emergence of specific morphologies and cell fates
- Acquisition of germ layer cell fates mimics early human embryonic diversity

## Introduction

In embryonic development, symmetry breaking events characterize the repeated emergence of different cell fates, which self-assemble into complex embryonic tissues via coordinated changes in cell adhesion. To this end, key three-dimensional states in which specific adhesions are relevant have been examined in work with model organisms and limited studies of human embryos. During compaction, the embryo forms a cystic cavity adjoined by the inner cell mass: a nonpolar group of cells that highly express the cell-cell adhesion molecule E-cadherin (CDH1) (1–5). The embryo then undergoes zona pellucida hatching, which involves epithelial sheet protrusion and subsequent expulsion of the early epiblast from its casing. Finally, during gastrulation, CDH1 is sharply downregulated and replaced by N-cadherin (CDH2), resulting in delamination and invasion of primitive streak into the space between the epiblast and developing yolk sac. Dysregulation of symmetry breaking during these key transitional periods often results in congenital disease or embryonic lethality, highlighting the importance of attaining a more mechanistic understanding of morphogenesis in the earliest stages of development (6, 7). Despite extensive research into how intercellular adhesion molecules regulate the movement of cell populations, the connection between changes in intercellular adhesions and lineage fate decisions has remained poorly characterized. Furthermore, studies of human embryos *ex vivo* have been limited, and the dynamics of human development have been difficult to interrogate *in vivo* due to the physical restrictions, optical opacity, and complex signaling milieu inherent to the developing embryo. Therefore, to study how human specific symmetry breaking events direct morphogenesis, it is essential to establish *in vitro* human systems that promote the coincident development of analogous heterogeneous cell populations. Human pluripotent stem cells (hPSCs) provide an unlimited source of cells that can mimic developmental differentiation processes and maintain the ability to self-organize into tissue-like structures, such as optic cups, gut organoids, or stratified cortical tissues (8–11). However, these models manifest inherently stochastic and un-reproducible differentiation (12, 13), limiting our understanding of the mechanisms that control and coordinate human morphogenesis. Therefore, new approaches to reliably control the emergence and organization of multiple germ cell types would greatly advance tissue modeling and organ developmental studies.

Recent studies have demonstrated that mechanical signals are important regulators of cell fate, and have implicated CDH1 not only as a regulator of cell-cell adhesions and sorting, but as a common transducer of external mechanical forces which contribute to mesendoderm versus ectoderm fate decisions (14, 15). In this study, we interrogate the interplay between mechanics and cell-cell adhesions in directing symmetry breaking events in 3D. Specifically, we employ hydrogel encapsulation to mimic the physically confined environment of the zona pellucida and induce heterogenous changes in adhesion through mosaic knockdown of CDH1 using CRISPRi in small cell aggregates. We show that hiPSC aggregates undergo changes in 3D structure as well as population emergence reminiscent of the three germ lineages of pre-implantation embryos. Additionally, the combination of CDH1 knockdown and encapsulation leads to specific emergence of extraembryonic populations in aggregates after 6 days of culture, highlighting potential regulation of trophoblast development triggered by the physical microenvironment. Recent studies demonstrating that mechanical signals are important regulators of cell fate have implicated CDH1 not only as a regulator of cell-cell adhesions and sorting, but as a common transducer of external mechanical forces that contribute to mesendoderm vs. ectoderm fate decisions (14–16). In this study, we interrogate the interplay between mechanics and cell-cell adhesions in directing symmetry breaking events in 3D pluripotent human cell aggregates. Specifically, we employ hydrogel encapsulation to mimic the physically confined environment of the zona pellucida and induce heterogenous changes in adhesion through mosaic knockdown of CDH1 using CRISPRi in hiPSC aggregates. We show that the aggregates undergo changes in 3D structure as well as population emergence of the three germ lineages similar to preimplantation embryos. Additionally, the specific combination of CDH1 knockdown and encapsulation leads to emergence of extraembryonic populations in aggregates after 6 days of culture, highlighting potential regulation of trophoblast development by integrated microenvironmental factors.

## Results

### Loss of CDH1 Promotes Protrusion Morphology when Encapsulated

To mimic the size-scale and environment of the pre-implantation blastocyst, 50-cell human hiPSC aggregates were encapsulated in 1.5% alginate beads mixed with laminin and cultured for 6 days. Alginate encapsulation served as a proxy for zona pellucida encapsulation by providing both a physical barrier (unfunctionalized alginate) and a signaling competent ECM (laminin) (Figure 1A). To model CDH1 loss during gastrulation, CDH1 knockdown was induced in either 0%, 25% or 100% of the aggregate population using a previously established CRISPRi system, where knockdown is triggered by doxycycline inducible production of dead Cas9-KRAB (14, 17). Post aggregation, both encapsulated and unencapsulated CDH1(+), CDH1(-), and mosaic aggregates displayed polarized behavior, creating a single layer of cells surrounding a cystic cavity in the center (Figure 1B). Over time, a bilayer of cells in CDH1(+) aggregates assembled at the surface of the growing aggregate and the central cyst remained intact, indicating that the polarity of human iPSCs in monolayer culture is initially maintained in suspension culture.

**Figure 1:**
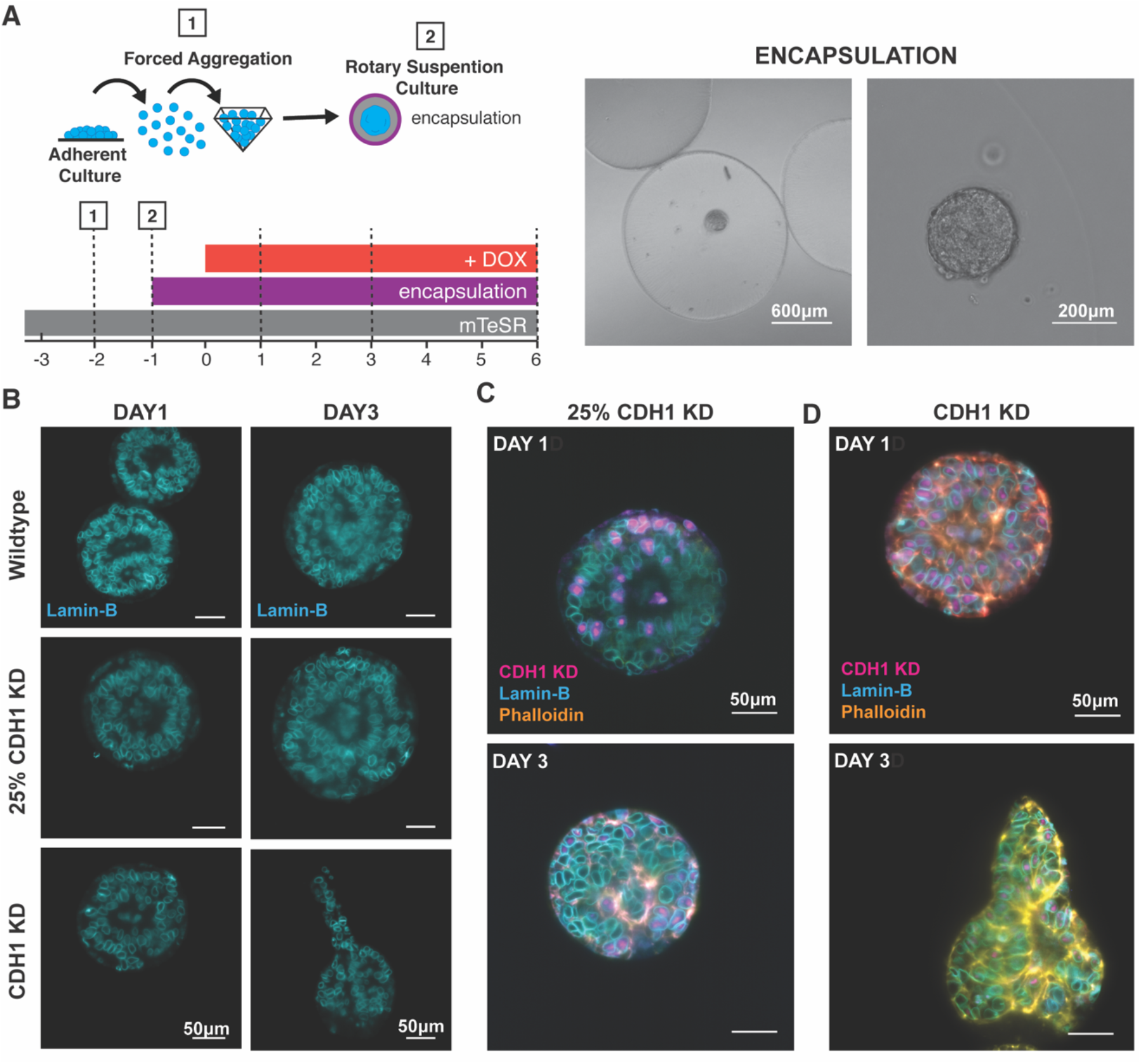
Loss of CDH1 promotes protrusion morphology. **(A)** A schematic of the experimental set up of encapsulation (left) and example images of encapsulated 50 cell human iPSC aggregates in alginate gel 20 minutes post encapsulation. **(B)** Optical sections of aggregates demonstrating evolution of morphologies on day 1 and day 3. **(C)** Optical section of mixed aggregates with 25% CDH1 knockdown. **(D)** Optical sections of aggregates lacking CDH1.

At day 3 of CDH1 knockdown, all aggregates consisted of bilayers surrounding a cystic cavity. In mixed aggregates, CDH1(+) and CDH1(-) cells physically sorted away from one another. The CDH1(-) cells clustered within the outer layer of the cystic aggregates, recapitulating the same spatial organization of cells engineered to express either high or low levels of CDH1 (18). Taken together, our results suggest that modulation of cell-cell adhesions alone is sufficient to induce radial organization of cell subtypes in hPSC aggregates (Figure 1C). Interestingly, while the CDH1(+) aggregates and the mixed aggregates retained a spherical shape, the CDH1(-) aggregates formed oblong protrusions extending away from the main body of the aggregate that reflected collective migration characteristics (Figure 1B,D). These extensions displayed similar morphologies to those present during zona pellucida hatching (1, 19, 20), in that extensions remained epithelial and no single cells were observed to migrate away from the aggregate into the alginate hydrogel material.

### Encapsulation Affects the Morphology of CDH1(-) Aggregates

To determine whether the observed extensions were a result of CDH1 knockdown or a response to encapsulation itself, human iPSC aggregates (50 cells each) were cultured in round bottom wells for 6 days without added extracellular matrix or alginate encapsulation (Figure 2A,B). Similar to encapsulated aggregates, cystic cavities formed in unencapsulated aggregates by day 1 persisted throughout culture. In mixed aggregates after 3 days, CDH1(+) cells segregated from CDH1(-) cells, mirroring the behavior seen in encapsulated mixed aggregates (Figure 2B, arrows mark segregated CDH1(-) cells). By day 5, CDH1(-) aggregates underwent a sheet delamination event characterized by an outer layer of mesenchymal-like cells peeling away from an inner epithelial core, displaying a cystic cavity (Figure 2C,D). Moreover, in the CDH1(-) condition, the entire aggregate lost OCT4 expression (Figure S1), the outer layer displayed EOMES expression, and the inner layer displayed SOX2 expression (Figure 2D), indicating a loss of pluripotency and subsequent divergence of cell populations into mesendoderm and ectoderm lineages, respectively. This lineage emergence is in contrast to the CDH1(+) aggregates which remained largely pluripotent (OCT4+SOX2+) or ectodermal (SOX2+) (Figure S1). These results suggest that silencing of CDH1 accelerates the loss of OCT4 in human iPSCs while still allowing for emergence of the mesendoderm and ectoderm lineages.

**Figure 2:**
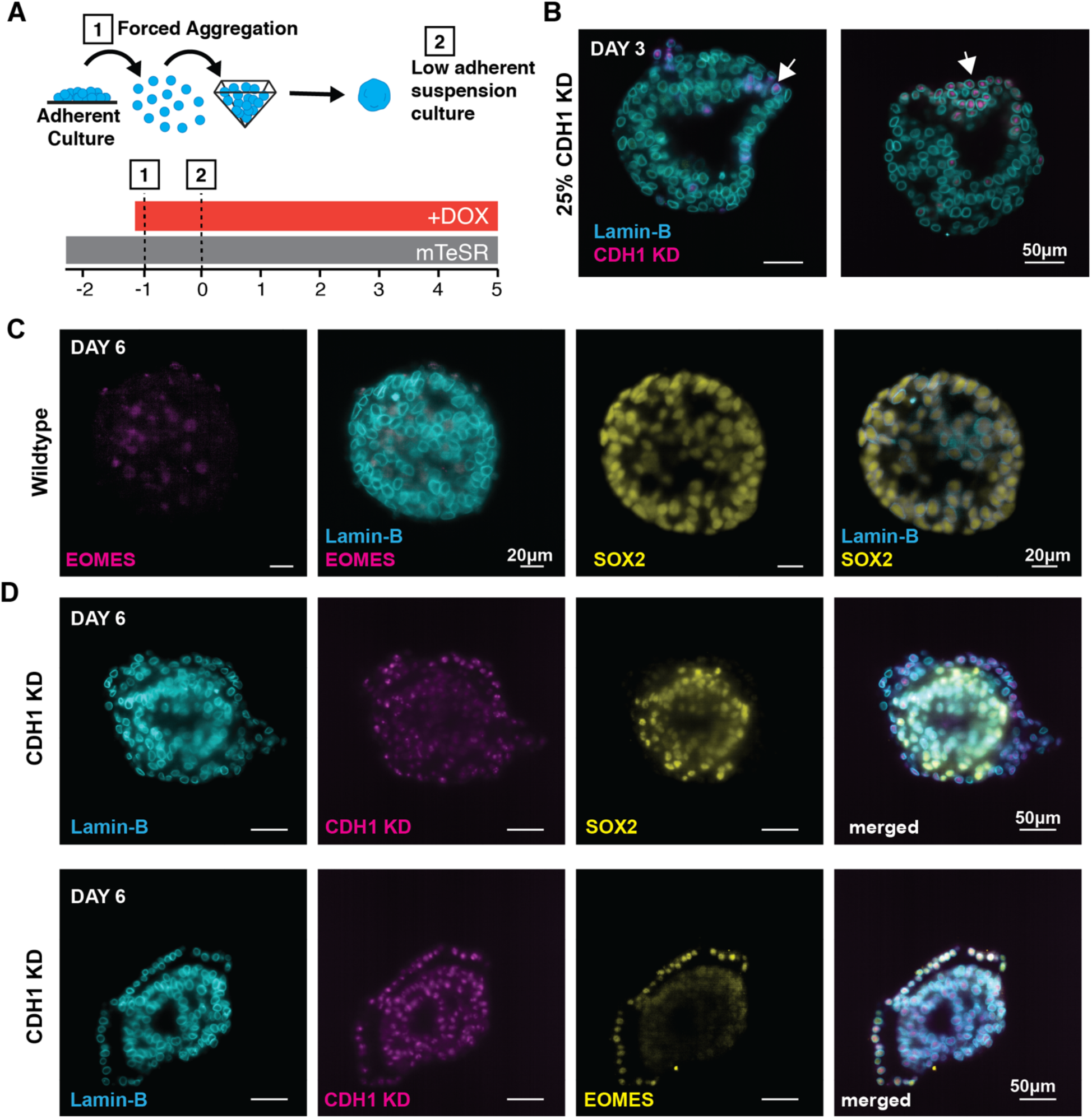
Encapsulation affects the morphology of CDH1(-) aggregates. **(A)** A schematic of the experimental set up. **(B)** Optical sections of mixed aggregates with 25% CDH1 knockdown demonstrating maintenance of cysts and segregation of CDH1(-) cells by day 3. **(C)** Optical sections of wildtype unencapsulated aggregates stained for EMOES and SOX2. **(D)** Optical sections of aggregates lacking CDH1 demonstrating changes in morphology and lineage fate spatial segregation.

### Emergence of Extraembryonic-like Cell Fates in Encapsulated Aggregates

Due to the marked differences in the morphology and cell type emergence between encapsulated/unencapsulated CDH1(+)/CDH1(-) aggregates, we used single-cell RNA sequencing to interrogate the diversity of cellular populations generated within the differentiating aggregates. The transcriptomes of encapsulated and unencapsulated aggregates at days 1, 3, and 6 post-aggregation were examined between CDH1(+), CDH1(-), and mixed aggregates (Figure 3A,B). The resulting clustered data set of single cell transcriptomes represented 8 cell states by lineage markers. At day 1, all aggregate types overlapped in a cluster marked by high expression of pluripotency genes (OCT4, SOX2, NANOG) (Figure 3C), whereas after 3 and 5 days aggregates transitioned through multiple lineage states, eventually clustering into populations expressing markers of mesendoderm, ectoderm, endoderm, and mesoderm (Figure 3C). Interestingly, clusters 8 and 10 displayed markers of the extraembryonic trophectoderm (CDX2, GATA3, CDH3, HAND1) (Figure 3C). Gene Ontology analysis of clusters 8 and 10 revealed gene networks associated with placental development and reproductive system development (Figure 3D). Furthermore, these clusters were largely populated by CDH1(-) cells that were encapsulated (Figure 3E), and contained highly expressed genes in pathways associated with trophectoderm development, such as Hippo signaling and non-canonical Wnt signaling (2, 21–23) (Figure 3F).

**Figure 3:**
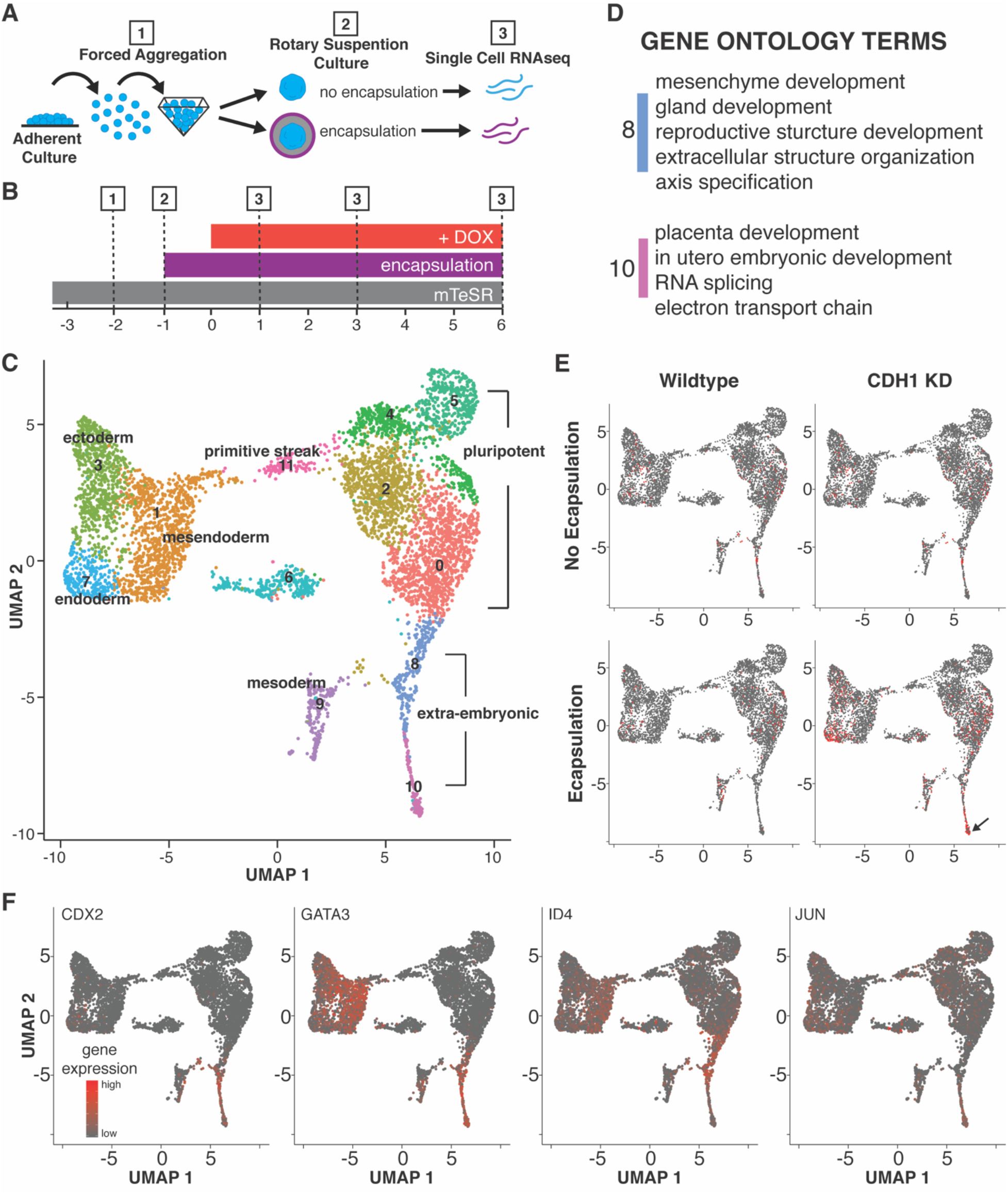
Single cell sequencing analysis of encapsulated and unencapsulated aggregates. **(A,B)** A schematic of the experimental set up. **(C)** UMAP demonstrating 12 clusters of cell populations at day 1,3, and 6. **(D)** Gene ontology terms for extraembryonic-like clusters 8 and 10 (p < 0.05). **(E)** Distribution of encapsulated and CDH1 knockdown cells within the UMAP clusters of all samples. **(F)** Distribution of genes associated with trophectoderm.

To computationally investigate whether this extraembryonic-like population (clusters 8 and 10) diverged from the other three germ lineages early on in the spontaneous aggregate differentiation, we examined cellular trajectories indicating transitions between states. Cellular trajectory reconstruction revealed that the differentiating population of encapsulated and nonencapsulated aggregates as a whole bifurcated after day 1, with one branch transitioning through primitive streak and then mesoderm lineage states, while a separate trajectory proceeded toward an extra-embryonic lineage state (Figure S1A). The mesendoderm, endoderm, and ectoderm germ lineages clustered separately from the lineage trajectories of pluripotency to mesoderm or to extraembryonic lineages (Figure S1A), indicating that the transition away from pluripotency may occur on a faster time scale that was not captured by sampling at days 1,3, and 5, and therefore preventing computational reconstruction of the complete lineage trajectory from pluripotency. This rapid time scale of differentiation is consistent with previous reports of fast switch-like exit from pluripotency (24), meaning that in order to capture lineage trajectories of cells as they lose pluripotency, more frequent sampling must occur. Overall, the early bifurcation between the germ lineages specific to the epiblast (mesoderm, endoderm, ectoderm) and the extraembryonic-like lineage in pseudo-time resembles the divergent population emergence at compaction of the *in vivo* embryo (Figure S1B), suggesting that this 3D human iPSC-based system can be used to model early developmental lineage diversification.

### Human iPSC Aggregates Have Similar Transcriptomes to Pre-implantation Human Embryos

To interrogate whether the emergence of lineages in our encapsulated and unencapsulated cell aggregates resembled that of the early human embryo (Figure 4A), our *in vitro* data set was directly contrasted with three previously published single cell transcriptome data sets from preimplantation embryos (25–27). *In vitro* aggregate cell transcriptomes were coclustered with single-cell RNA sequencing of preimplantation embryo stages from oocyte through late blastocyst from Yan *et al*. (Figure 4B). Embryonic stages from the oocyte to 4-cells clustered far from our *in vitro* aggregates (Figure 4B, red arrows), with only 33% clustering overlap between 4-cell embryos and 3D hiPSC aggregates, reflecting the lack of totipotent cell-like transcriptional profiles *in vitro*. Cells from the morula and late blastocyst co-clustered with the embryonic-like fractions of *in vitro* aggregates (Figure 4B, blue arrow heads) including in regions of primitive streak and germ-lineage restricted cells, demonstrating that *in vitro* aggregates reflect similar lineage transitions to those which occur *in vivo*. Interestingly, cells from 8-cell and morula stages had similar transcriptomes to *in vitro* extraembryonic-like cells from the encapsulated aggregates, marked by high CDX2 and HAND1 expression, possibly reflecting the *in vivo* transition from totipotency towards lineage-restricted embryonic and extra-embryonic cell types. Overall, coclustering with the Yan *et al*. dataset indicated that *in vitro* aggregate cell transcriptomes recapitulate aspects of pre-implantation embryos from approximately the 8-cell stage through late blastocyst, with the majority of cells reflecting a late-blastocyst-like phenotype.

**Figure 4:**
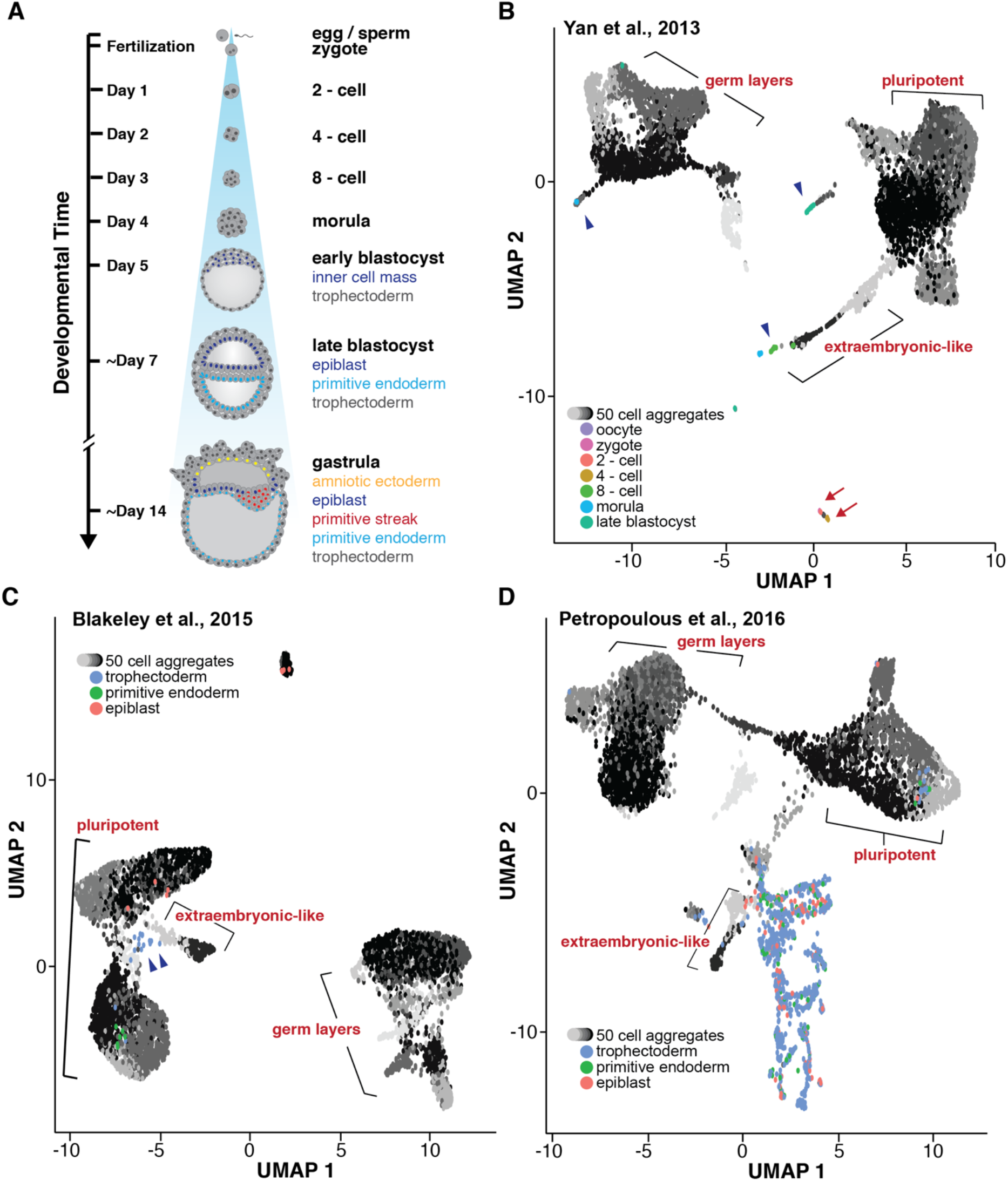
Comparison of *in vitro* aggregate transcriptome with preimplantation human embryos. **(A)** Schematic of early human embryonic development. **(B-D)** UMAPs displaying 50 cell encapsulated and non-encapsulated aggregate transcriptomes (gray) with three single-cell sequencing data sets (Yan *et al*., Blakeley *et al*., and Petropoulous *et al*., respectively) where the extraembryonic-like population is labeled with brackets.

To confirm the approximate staging of *in vitro* aggregates with the late blastocyst, the transcriptional profiles were subsequently aligned with two independent single cell sequencing data sets of late blastocyst embryos (Figure 4C,D). In both Blakeley *et al*. and Petropoulous *et al*., cells from the trophectoderm clustered close to the extraembryonic-like fraction, confirming that *in vitro* aggregates contain a trophectoderm-like population of cells. There was 100% alignment overlap between cells from the primitive endoderm and epiblast in the Blakeley *et al*. data set and the embryonic-like fraction of *in vitro* aggregates, with epiblast cells aligning better to the pluripotent and transitioning fractions, while primitive endoderm cells aligned more with *in vitro* cells in the differentiating compartment (Figure 4C). Additionally, 18.8% of the trophectoderm cell transcriptomes clustered with the extra-embryonic like compartment of cells from the *in vitro* aggregates (Figure 4C, blue arrow heads). Alignment with primitive endoderm and epiblast cells was poorer in the Petropoulous *et al*. data set, possibly reflecting differences in the developmental stage of the *in vitro* and *in vivo* cells, where *in vitro* iPSCs presumably reflect a population of cells in a primed pluripotency state whereas the *in vivo* population of cells in the epiblast would represent a more naïve state of pluripotency. Despite this reduced overlap between populations, 36.1% of primitive endoderm cells and 28.9% of epiblast cells still aligned with the differentiating compartment of the *in vitro* aggregates, again suggesting that *in vitro* aggregates recapitulate the developmental trajectories of cells in the developing embryo (Figure 4D). Overall, the transcriptomic similarities between our *in vitro* results and the three human embryo data sets suggest that hydrogel encapsulation coupled with CDH1 knockdown enhances activity of gene regulatory networks that mimic developmental processes in pre-implantation embryos.

## Discussion

The emergence and coincident organization of multiple lineages in the developing embryo is essential for the formation of the overall body plan and the development of functional tissues. However, the process by which cell-cell adhesions, ECM/mechanical signals, and morphogen cues coordinate to direct symmetry breaking and subsequent tissue formation *in vivo* is not well understood. In this study, we demonstrate that culture of individual encapsulated human iPSC aggregates within hydrogel environments yields aspects of early human development and lineage emergence, even in the absence of adding any exogenous morphogens. Additionally, attenuation of CDH1 within encapsulated aggregates enabled the emergence of an extraembryonic-like population of cells reminiscent of trophoblast cells. These results demonstrate that the physical structure of iPSC culture is sufficient to initiate symmetry breaking, and that adhesion changes in the developing embryo may directly control lineage decisions, versus simply emerging as a result of fate commitment. Furthermore, this study suggests that the physical microenvironment plays a valuable role in the coordination of lineage fate decisions, highlighting a possible regulatory mechanism of both adhesions and mechanical cues employed by the early embryo to control population emergence.

As the embryo develops, it undergoes a series of cell polarity changes, where cells are segregated into an apical and basal domain, that dictate developmental transitions. For example, compaction relies on the polarization of the trophoblasts to allow for cavity formation (2, 23), while gastrulation relies on the loss of polarity as invaginating cells begin their migration across the embryo As the embryo develops it goes through regimented changes in polarity that dictate developmental transitions. For example, compaction relies on the polarization of the trophoblasts to allow for cavity formation (2, 23), whereas gastrulation relies on the loss of polarity as invaginating cells begin their migration across the embryo (21,28). This previous work in combination with our studies suggest that the loss of adhesions in aggregates due to CDH1 knockdown triggers a signaling network that regulates polarity-dependent lineage transitions.

The mechanical microenvironment has been shown to regulate lineage fate emergence in pluripotent stems cells *in vitro*, suggesting that a similar mechanism may regulate embryonic cell fate (15,29, 30). The emergence of a trophoblast-like population, more so in encapsulated CDH1 knockdown aggregates, suggests that the physical microenvironment provided by encapsulation in tandem with changing surface adhesion properties provides necessary cues for extraembryonic fate specification. Trophoblast development is primarily regulated by Hippo signaling, a well-established mechanically responsive signaling pathway (22, 31). Additionally, it has been previously reported that the mechanical micro-environment has the potential to regulate lineage fate emergence in pluripotent stems cells, suggesting that a similar mechanism *in vitro* may regulate embryonic cell fate (15,29,30). The emergence of a trophoblast-like population, particularly in encapsulated CDH1 knockdown aggregates, suggests that the physical microenvironment provided by encapsulation provides necessary cues for extraembryonic fate specification. Interestingly, trophoblast development is primarily regulated by Hippo signaling, a mechanically responsive signaling pathway (22,31). Thus, it is possible that the enhanced emergence of trophoblast-like cells with encapsulation is a result of a mechanically triggered upregulation in Hippo signaling.

Overall, this study demonstrates how physical environmental parameters and intercellular adhesive properties of iPSCs can cooperatively impact developmental processes that regulate 3D morphogenesis and lineage emergence in the early embryo. Our results demonstrate that the combinatorial effect of adhesion regulation and microenvironment impacts lineage emergence and population morphogenesis in human iPSCs, potentially reflecting mechanisms within the early human embryo that robustly regulate development. Ultimately, this work provides insights that are relevant to stem cell biology and human embryonic development, facilitating routes to improve the robustness of *in vitro* differentiation methods as well as potentially illuminating strategies relevant for therapeutics, such as *in vitro* fertilization.

## Experimental Procedures

### Human Induced Pluripotent Stem Cell Culture

All work with human induced pluripotent stem cells (iPSCs) was approved by the University of California, San Francisco Human Gamete, Embryo, and Stem Cell Research (GESCR) Committee. Human iPSC lines were derived from the Allen Institute WTC11-LaminB cell line (AICS-0013 cl.210). All cell lines were karyotyped by Cell Line Genetics and by qPCR and reported to be karyotypically normal. Additionally, all cell lines tested negative for mycoplasma using a MycoAlert Mycoplasma Detection Kit (Lonza).

Human iPSCs were cultured on growth factor reduced Matrigel (Corning Life Sciences) and fed daily with mTeSR™-1 medium (STEMCELL Technologies) (32). Cells were passaged by dissociation with Accutase (STEM CELL Technologies) and re-seeded in mTeSR™-1 medium supplemented with the small molecule Rho-associated coiled-coil kinase (ROCK) inhibitor Y-276932 (10 μM; Selleckchem)(33) at a seeding density of 12,000 cell per cm^2^.

The CRISPRi CDH1 cell line was generated via insertion of a previously published Tet-ON system inserted into the AAVS1 locus via TALENS (Addgene plasmid # 73498; Mandegar et al., 2016) into the AICS-0013 cl.210 parent line. The previously published guide RNA sequence used to target CDH1 (GCAGTTCCGACGCCACTGAG) was cloned into the gRNA expression vector (addgene plasmid # 73501) using a BsmBI restriction enzyme cloning strategy described in Mandegar et al. The guide RNA vector was then electroporated into the parent line containing the CRISPRi system using the Human Stem Cell Nucleofector Kit 1 solution with the Amaxa nucleofector 2b device (Lonza). Nucleofected cells were then seeded into a 6-well plate in mTeSR™-1 supplemented with Y-27632 (10 μM) and underwent blasticidin (ThermoFisher Scientific; 10 μg/ml) selection for 6 days. Surviving cells were then colony picked, expanded, and knockdown tested via qPCR after 5 days CRISPRi induction via addition of doxycycline (2μM) to the culture media (Figure S2).

### Encapsulated Mixed Aggregate Generation

Cell aggregates of ~50 cells were created using 400 × 400 μm PDMS microwell inserts in 24-well plates (~975 microwells per well), similar to previously published protocols (34–36). Dissociated iPSC cultures were resuspended in mTeSR™-1 supplemented with Y-27632(10μM), mixed at proper ratios and concentration (50 cells/well), added to microwells, and centrifuged (200 rcf). After 18 hours of formation, 50 cell aggregates were transferred into 1.5% ultrapure MVG alginate (Pronova) mixed with Laminin from Engelbreth-Holm-Swarm murine sarcoma (6μg/mL; Sigma Aldrich) at a concentration of 16,000 aggregates/mL alginate. Alginate solution was prepared by mixing the appropriate amount of alginate to generate a 1.5% solution into calcium-free DMEM (Gibco) and sterilized by autoclave. Beads encapsulating single aggregates were generated using an electrostatic bead generator (Nisco). A 400μm nozzle and syringe pump (flow of 6mL/hour) was used to extrude alginate solution with aggregates and dropped into a 100 mM calcium chloride (EMD) bath to trigger hardening of the alginate into a gel. Encapsulated aggregates were then washed 3X with DPBS containing calcium and magnesium (ThermoFisher Scientific) and once with mTeSR™-1 medium (STEMCELL Technologies). Encapsulated aggregates were allowed to recover for 24 hours in mTeSR™-1 medium (STEMCELL Technologies) in rotary suspension, then fed daily with mTeSR™-1 medium supplemented with doxycycline (DOX)(2μM) to induce CDH1 knockdown.

### Un-encapsulated Mixed Aggregate Generation

Unencapsulated 50 cell aggregates were created by dissociation of human iPSCs with Accutase (STEM CELL Technologies) and reseeding in mTeSR™-1 medium (STEMCELL Technologies) supplemented with Y-276932 (10μM; Selleckchem) into 96-well non-adherent round bottom plates, with a total of ~50 cells were seeded per well. After 18 hours of aggregate formation, Y-276932 was removed from the media and DOX(2μM) was supplemented into the mTeSR™-1 medium to induce CDH1 knockdown. Aggregates were then fed daily with mTeSR™-1 medium supplemented with DOX (2μM).

### Immunofluorescence Staining and Imaging

Aggregates were unencapsulated by washing 3X with a sodium citrate solution (55mM, Sigma), fixed with 4% paraformaldehyde (VWR) for 40 minutes, and then washed three times with PBS. Aggregates to be used for histology were embedded in HistoGel Specimen Processing Gel (Thermo Fisher) prior to paraffin processing. Paraffin embedded samples were sectioned in 5μm sections, baked for 1 hour at 60°C, and subsequently stained for H&E. For immunofluorescent staining, epitope retrieval was performed by submersing slides in Citrate Buffer pH 6.0 (Vector Laboratories) in a 95°C water bath for 35min. Samples were permeabilized in 0.2% Triton X-100 (Sigma-Aldrich) for 5min, blocked in 1.5% normal donkey serum (Jackson Immunoresearch) for 1hour, and probed with primary and secondary antibodies against SOX2, PAX6, T, NES, TUBB3, and CDH2 (Table S3). Coverslips were mounted with anti-fade mounting medium (ProlongGold, Life Technologies) and samples were imaged on a Zeiss Axio Observer Z1 inverted microscope equipped with a Hamamatsu ORCA-Flash 4.0 camera.

### Whole Mount Lightsheet Imaging

4% paraformaldehyde-fixed paraffin-embedded samples (see “*Histology, Immunocytochemistry, and Imaging*”) were permeabilized with 0.3% Triton X-100 (Sigma-Aldrich) for 5min, blocked in 5% normal donkey serum (Jackson Immunoresearch) for 1 hour, and probed with primary and secondary antibodies against PAX6 and T (TableS3) for 2 hours. Samples were then embedded in 1.5% low melt agarose (BioReagent) and drawn up into ~1mm diameter imaging capillaries and subsequently imaged on the Zeiss Z.1 Light sheet Microscope equipped with a PCO.edge SCMOS camera.

### Single Cell RNA Sequencing Sample and Library Preparation

Multiple organoid samples were combined and processed together using the MULTI-Seq technology (37). Organoids were singularized using Accutase (STEMCELL Technologies) and washed with cold PBS. Cells were resuspended in PBS with lipid-modified Anchor and Barcode oligonucleotides (kindly provided by Dr. Zev Gartner) and incubated on ice for 5 min. A co-Anchor oligo was then added in order to stabilize membrane retention of the barcodes incubated for an additional 5 min on ice. Excess lipid-modified oligos were quenched with 1 % BSA in PBS, washed with cold 1% BSA solution, and counted using a Countess II FL (Life Technologies). Single cell GEMs and subsequent libraries were then prepared using the 10X Genomics Single Cell V2 protocol with an additional anchor specific primer during cDNA amplification to enrich barcode sequences (37). Short barcode sequences (approx. 65-100bp determined by Bioanalyzer) were purified from cDNA libraries with two sequential SPRI bead cleanups. Barcode library preparation was performed according to the KAPA HiFi Hotstart (Kapa Biosystems) protocol to functionalize with the P5 sequencing adapter and library-specific RPIX barcode. Purified ~173bp barcode fragments were isolated with another SPRI bead cleanup and validation by Bioanalyzer.

The sample library was sequenced on an Illumina NovaSeq yielding an average of 41,112 reads per cell and 6,444 cells. The MULTI-Seq barcode library was sequenced on an Illumina NextSeq yielding an average of 9,882 reads per barcode and enabling sample assignment for 4,681 of 6,124 unique UMIs detected (76.4% recovery), using the demultiplexing code provided by the MULTI-Seq protocol (37).

### Genome Annotation, RNA-seq Read Mapping, and Estimation of Gene and Isoform Expression

The sample library was aligned to the human GRCh38 reference genome using Cell Ranger v1.2.0 (10x Genomics). Gene expression levels were assessed using the Seurat v3.0.0 analysis pipeline (38). First cells with fewer than 200 detected genes, fewer than 1,000 total detected transcripts, or greater than 10% mitochondrial gene expression were removed. Next, expression levels were log normalized, and the top 2,000 variable genes calculated using the VST algorithm. The top 20 principal components were used to group cells into 12 clusters using a resolution of 0.4. Finally, top markers were detected for each cluster by detecting the top differentially expressed genes between one cluster and the remaining data set, where at least 25% of cells in the cluster expressed the gene and the gene was expressed at least 0.25 log2 fold-change different from the remaining population. Clusters and gene expression were visualized on a two-dimensional UMAP projection of the first 20 principal components.

### Data Acquisition, Processing, and Merging

Human single cell RNA sequencing data sets from previously published papers (25–27) were downloaded from Gene Expression Omnibus (GEO66507, GSE36552, E-MTAB-3929). FastQ data sets were aligned to the human GRCh38 reference genome using STAR aligner (39) for GEO66507 and GSE36552 and using TopHat2 (40) for E-MTAB-3929 prior to generating a counts matrix using the FeatureCounts software (41). For the entire data set, all counts matrices were concatenated into one matrix. Each matrix was then read as a Seurat object, which could then be combined with other data sets and analyzed using the Seurat v3.0.0 analysis pipeline (38). Batch correction was performed using the Harmony algorithm (42).

### Cluster Analysis

To assign cluster identity, the top markers for each cluster were tested for GO term enrichment using the biological process “enrichGO” function in the R package “clusterProfiler” v3.12. (43) In addition, differentiation and lineage specification in each cluster was assessed by examining expression level of panels of pluripotency, mesendoderm, endoderm, ectoderm, mesoderm, and trophectoderm markers. Computational trajectory analysis of transitions between cell states was performed using Monocle 3 (44).

## Acknowledgements

We would like to thank the Gladstone Light Microscopy and Histology Core, the Gladstone Stem Cell Core and The Gladstone Flowcytometry Core for their support and experimental expertise. Additionally, we would like to specifically thank Dr. Vaishaali Natarajan for her expertise and assistance with encapsulation and Dr. Zev Gartner and his lab for reagents and expertise in the MULTI-seq assay.

## Author Contributions

ARGL and TCM were responsible for the conceptualization of the project and designing experiments. Differentiations were conducted by ARGL and IV. Single cell RNA sequencing preparation and analysis were performed by ARGL, DAJ, and MZK. Image analysis script was written and implemented by DAJ and ARGL. Light sheet microscopy and analysis was conducted by ARGL, DAJ, and IV. Cell lines were generated by ARGL, DAJ, MZK, and NMC. ARGL and IV prepared the figures with input from all co-authors. All co-authors contributed to writing the original manuscript.

## Figure Legends

**Figure S1:**
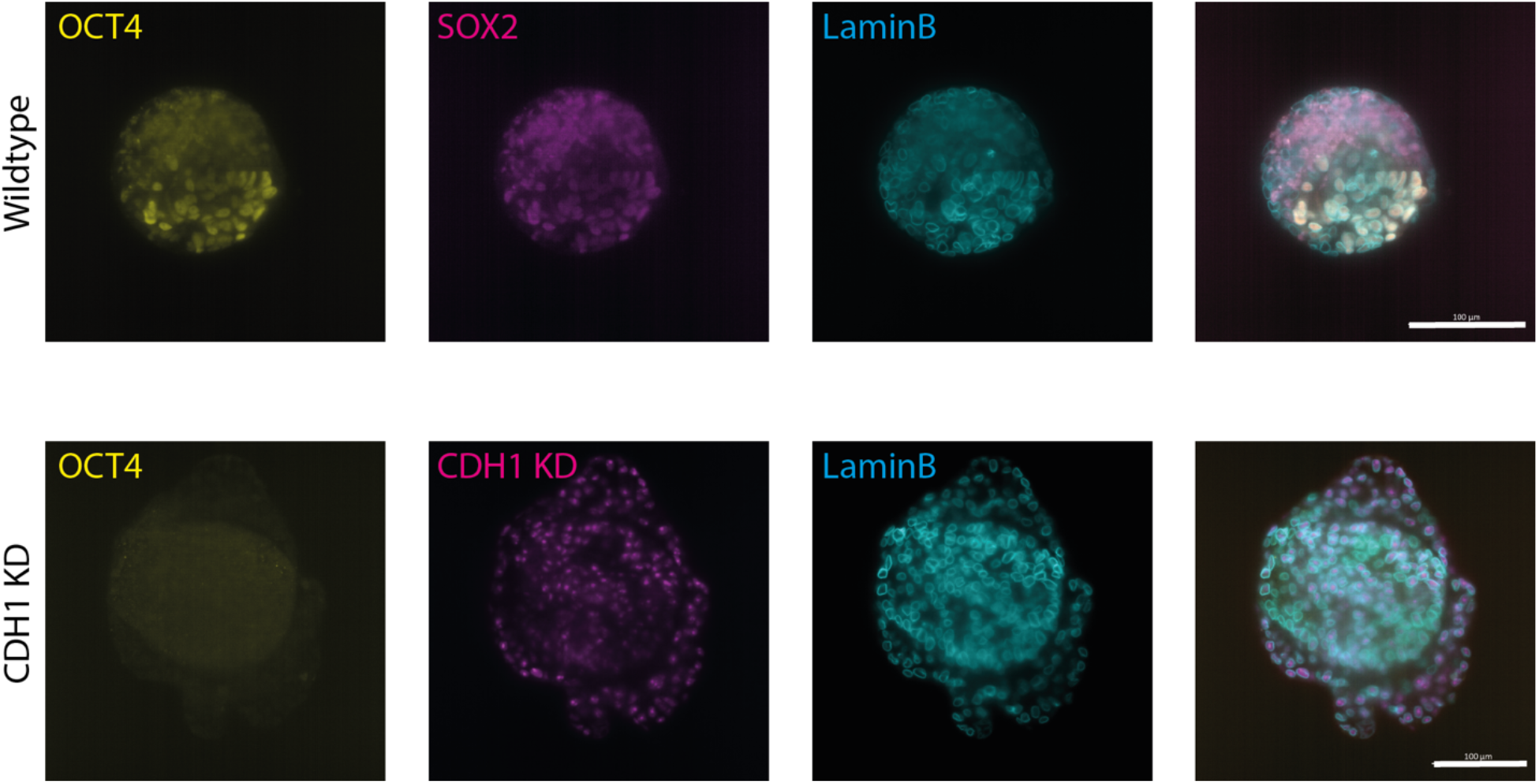
OCT4 staining in encapsulated aggregates. (A) Immunofluorescence images of OCT4 staining in encapsulated aggregates at day 5 (scale bars = 100um).

**Figure S2:**
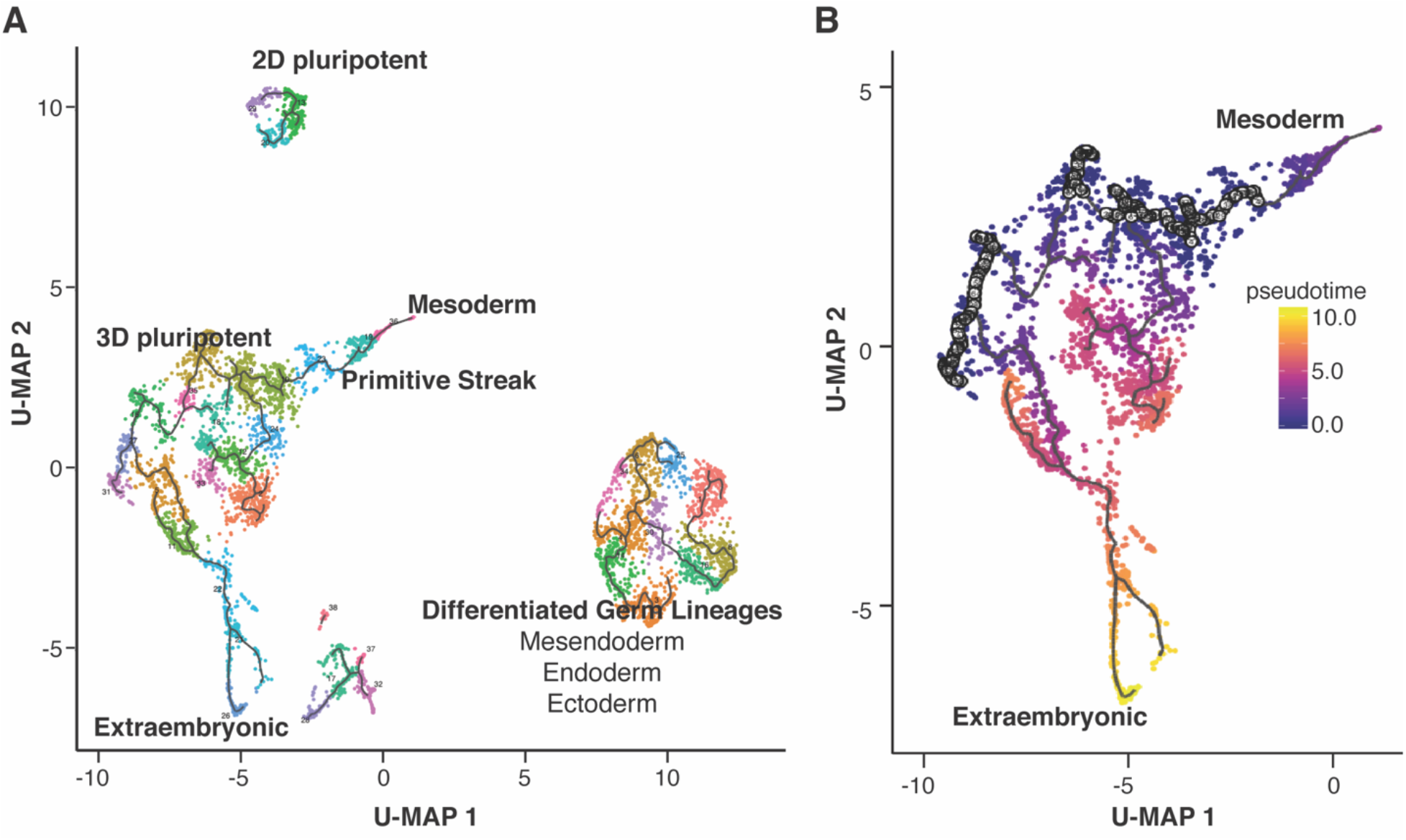
Lineage trajectories of 50 cell aggregates. **(A)** Lineage trajectory reconstruction using Monacle demonstrating Mesoderm and Extraembryonic cell fates on separate lineage branches. **(B)** Pseudotime reconstruction of the mesoderm and extraembryonic lineages.

**Figure S3:**
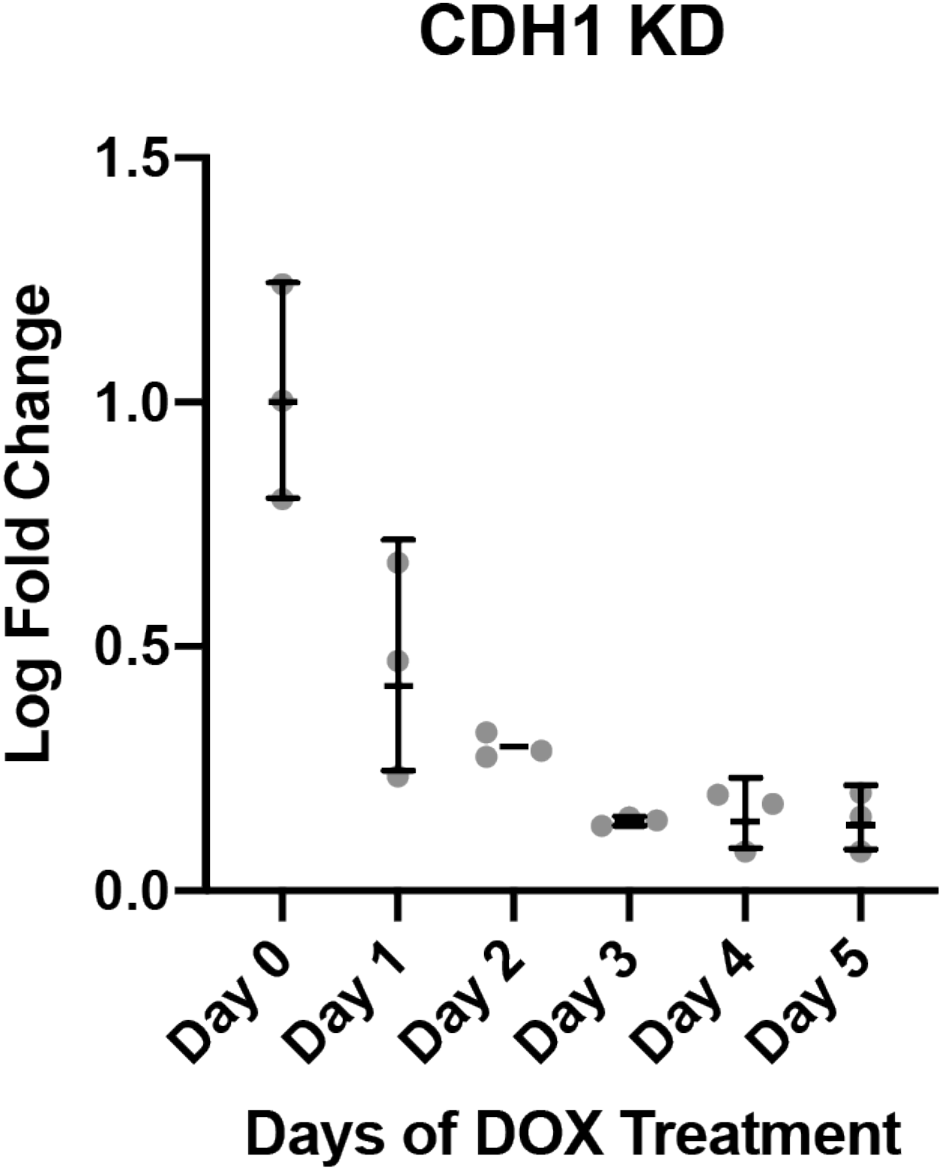
Karyotype and knockdown testing of CRISPRi CDH1 line. **(A)** qPCR plot of loss of CDH1 expression over a 5 day time course of DOX addition to cell culture media (n=3).

